# One Health genomic surveillance of *Escherichia coli* demonstrates distinct lineages and mobile genetic elements in isolates from humans versus livestock

**DOI:** 10.1101/434001

**Authors:** Catherine Ludden, Kathy E. Raven, Dorota Jamrozy, Theodore Gouliouris, Beth Blane, Francesc Coll, Marcus de Goffau, Plamena Naydenova, Carolyne Horner, Juan Hernandez-Garcia, Paul Wood, Nazreen Hadjirin, Milorad Radakovic, Nicholas M. Brown, Mark Holmes, Julian Parkhill, Sharon J. Peacock

## Abstract

Livestock have been proposed as a reservoir for drug-resistant *Escherichia coli* that infect humans. We isolated and sequenced 431 *E. coli* (including 155 ESBL-producing isolates) from cross-sectional surveys of livestock farms and retail meat in the East of England. These were compared with the genomes of 1517 *E. coli* associated with bloodstream infection in the United Kingdom. Phylogenetic core genome comparisons demonstrated that livestock and patient isolates were genetically distinct, indicating that *E. coli* causing serious human infection do not directly originate from livestock. By contrast, we observed highly related isolates from the same animal species on different farms. Analysis of accessory (variable) genomes identified a virulence cassette associated previously with cystitis and neonatal meningitis that was only present in isolates from humans. Screening all 1948 isolates for accessory genes encoding antibiotic resistance revealed 41 different genes present in variable proportions of humans and livestock isolates. We identified a low prevalence of shared antimicrobial resistance genes between livestock and humans based on analysis of mobile genetic elements and long-read sequencing. We conclude that in this setting, there was limited evidence to support the suggestion that antimicrobial resistant pathogens that cause serious infection in humans originate from livestock.

**Importance:** The increasing prevalence of *E. coli* bloodstream infections is a serious public health problem. We used genomic epidemiology in a One Health study conducted in the East of England to examine putative sources of *E. coli* associated with serious human disease. *E. coli* from 1517 patients with bloodstream infection were compared with 431 isolates from livestock farms and meat. Livestock-associated and bloodstream isolates were genetically distinct populations based on core genome and accessory genome analyses. Identical antimicrobial resistance genes were found in livestock and human isolates, but there was little overlap in the mobile elements carrying these genes. In addition, a virulence cassette found in humans isolates was not identified in any livestock-associated isolate. Our findings do not support the idea that *E. coli* causing invasive disease or their resistance genes are commonly acquired from livestock.

## INTRODUCTION

*Escherichia coli* is a leading cause of infection in hospitals and the community (1, 2). The prevalence of *E. coli* bloodstream infections has shown a marked increase in Europe and the United States since the early 2000’s (3–6). This has been associated with the emergence and global dissemination of *E. coli* that produce extended-spectrum β-lactamases (ESBL-*E. coli*). Such isolates are resistant to many penicillin and cephalosporin antibiotics (3, 5), and are associated with excess morbidity, mortality, longer hospital stay and higher healthcare costs compared with infections caused by *E. coli* that are not ESBL producers (7–9). Successful therapy has been further challenged by the emergence of multidrug-resistant (MDR) *E. coli* with acquired resistance to the carbapenem drugs and more recently to colistin, a drug of last resort for multidrug-resistant infections (10, 11).

Tackling the rising trends in prevalence of MDR *E. coli* infections in humans requires an understanding of reservoirs and sources for human acquisition. Food-producing animals have been proposed as a source of ESBL-*E. coli* in humans based on comparison of bacterial genotypes using multilocus sequence typing (MLST) (12–14). This method lacks sufficient discrimination to generate robust phylogenetic comparisons of population genetics and does not capture information on accessory genome composition such as genes encoding drug resistance. Whole genome sequencing overcomes both of these limitations but there are limited published data on the use of this technique to address the transmission of antibiotic resistance *E. coli* between livestock and humans (15). Here, we report the findings of a genomic epidemiological investigation of *E. coli* sourced from livestock, meat and patients with bloodstream infection within a tightly defined geographical location.

## RESULTS

### Isolation of *E. coli* from livestock farms and retail meat

A cross-sectional survey was performed between 2014 and 2015 to isolate ESBL-*E. coli* and non-ESBL producing *E. coli* from livestock at 29 farms in the East of England, United Kingdom (UK) (10 cattle (5 beef/5 dairy), 10 pig, and 9 poultry (4 chicken/5 turkey)) (Fig. 1b). A total of 136 pooled faecal samples (34 from cattle, 53 from pigs, 49 from poultry) were collected and cultured. *E. coli* was isolated from all 29 farms and ESBL-*E. coli* was isolated from 16 (55%) of these (Table S1). The highest prevalence of ESBL-*E. coli* occurred in poultry farms (8/9), followed by pig (5/10) and cattle farms (3/10).

**Fig. 1.**
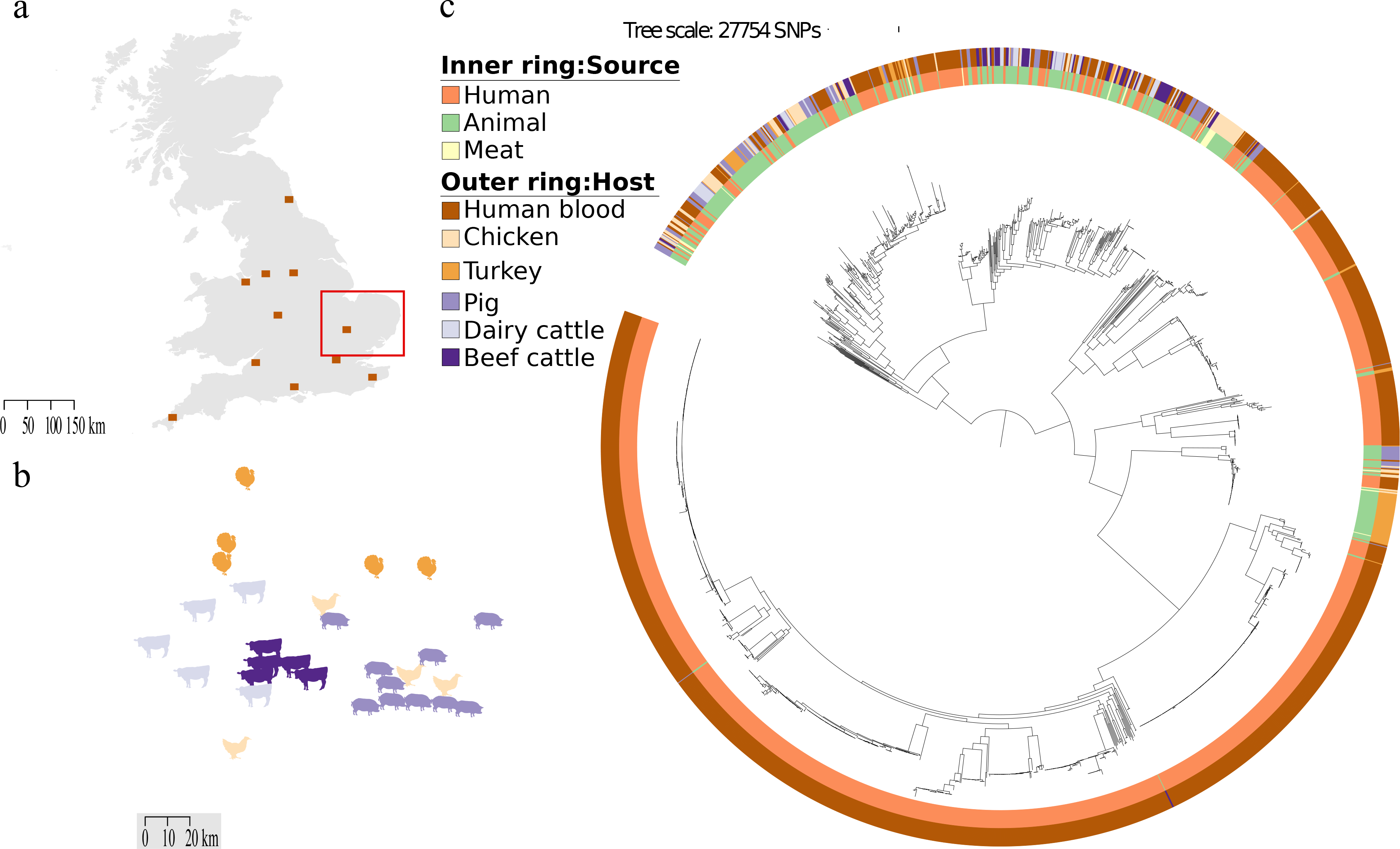
a: Map of the UK showing locations for the human clinical isolates, with the East Anglian region highlighted by a red box. b: Map of the East Anglian region showing the locations of farms (images indicate livestock species). c: Maximum likelihood tree based on SNPs in the genes core of 1948 *E. coli* isolates cultured from livestock farms, retail meat and patients with bloodstream infection.

A cross-sectional survey was performed in April 2015 to isolate ESBL-*E. coli* from 97 pre-packaged fresh meat products purchased in 11 major supermarkets in Cambridge, East of England (5-16 products per supermarket, see Table S2 in supplemental material). These originated from 11 different countries although the majority (69/97, 71%) were from the UK. ESBL-*E. coli* was isolated from 19/97 (20%) products, of which 16 were chicken originating from the UK (n=12), Ireland (n = 2), Hungary (packaged in Ireland, n=1) or multiple origins (Brazil, Thailand and Poland, n=1), and 3 were from turkey (n=2) and pork (n=1) from the UK.

We sequenced a total of 431 *E. coli* isolates from livestock (n=411) and meat (n=20), of which 155 were ESBL-*E. coli* (livestock=136, meat =19).

### Evaluation of the *E. coli* collection from patients with bloodstream infection

A key study objective was to determine whether livestock and retail meat represented potential sources of *E. coli* associated with serious invasive disease in humans. In light of this, the human isolates used in the comparison with livestock isolates had caused bloodstream infection. A total of 1517 open access *E. coli* genomes (ESBL=142, non-ESBL=1375) associated with bloodstream infection were retrieved (16, 17). Bloodstream isolates obtained from patients admitted to the Cambridge University Hospitals NHS Foundation Trust in the East of England between 2006 and 2012 (n=424) (16, 17) were combined with bloodstream isolates submitted to British Society of Antimicrobial Chemotherapy from 10 further hospitals across England and CUH (n=1093) between 2001 and 2011 (locations shown in Fig. 1a, and full isolate listing in Table S1) (16, 17). A potential limitation of this human isolate collection is that they might over-represent hospital-acquired isolates, whilst a comparison of *E. coli* from livestock would require a comparison of community-acquired bacteria. Two analyses were undertaken to evaluate this possibility. First, we defined where the bloodstream infection was acquired for 1303 cases for whom we had this information. This demonstrated that 886/1303 (66%) cases were community-associated. We then constructing a maximum likelihood tree of the invasive disease genomes to compare the phylogeny of isolates associated with community- versus healthcare-associated disease (Fig. S1). This demonstrated that genomes from the two categories were intermixed and distributed across the phylogeny, with no evidence of clustering by origin of infection. We concluded that our invasive collection was likely to include strong representation of *E. coli* carried by people in the community.

### Genomic comparison of *E. coli* from patients with bloodstream infection, livestock, and retail meat

We combined and compared the 1517 human invasive with the 431 livestock-associated genomes. Analysis of the 1948 genomes identified 331 multilocus sequence types (STs), 44 clonal complexes (CCs) and 149 singletons (STs that did not share alleles at six out of seven loci with any other STs). Most STs were host-specific, with 192 human-specific STs (1261/1517 isolates, 83%), 98 livestock-specific STs (225/411 isolates, 55%) and 4 meat-specific STs (4/20 isolates, 20%). Thirty-five STs contained isolates from both humans and livestock/meat (n=431), while 2 STs were only found in isolates from livestock and meat (n=27) (Table S1). The three most common STs associated with bloodstream infection were ST73, 131 and 95, whilst the three most common STs associated with livestock were ST10, 17 and 602 (Table S1). The greatest overlap in STs between the two reservoirs occurred in ST10 (3%, 16%, 0% of human, livestock and meat isolates, respectively), and ST117 (1%, 8%, 20% of human, livestock and meat isolates, respectively).

Phylogenies based on single nucleotide polymorphisms (SNPs) in the core (conserved) genomes of isolates representing CC10 (n=149) and CC117 (n=64) demonstrated that human isolates were intermixed with livestock isolates in CC10, but were generally distinct from livestock isolates in CC117 (Supplementary Figs. S2 & S3). Pairwise SNP analysis demonstrated that the most closely related human/livestock isolate pairs were 85 and 96 SNPs different for CC10 and CC117, respectively. The estimated mutation rate for *E. coli* is one SNP/core genome/year (18, 19), and so CC10 and CC117 isolates in humans and livestock was not associated with recent transmission between the two groups. Combining the study CC117 isolates with 7 publicly available ST117 genomes (ERR769196, ERR769195, ERR769183, ERR769169, SRR1314275, SRR3410778 and SRR3438297) in a Bayesian phylogenetic analysis provided further evidence for the lack of recent transmission between human and livestock hosts in our study. The dated phylogeny revealed a UK cluster of 47 CC117 isolates (containing 44 turkey, 1 chicken and 2 human isolates), for which the estimated time of most recent common ancestor (TMRCA) was 1989 (95% highest posterior density interval [HPD] 1979-1996), coinciding with the first global report of *bla*_CTX-M-1_ (20). Of the 47 isolates, 36 (77%) carried *bla*_CTX-M-1_, which was uncommon in the rest of the bacterial population. All 36 isolates were from turkeys, representing a *bla*_CTX-M-1_ poultry-associated lineage, for which the TMRCA was 2011 (95% HPD 2010-2013) (Fig. S4), suggesting acquisition of *bla*_CTX-M-1_ by this lineage between 1989 and 2011.

We then compared the genetic relatedness of the 431 livestock/meat *E. coli* with the 1517 *E. coli* associated with human bloodstream infection. A maximum-likelihood phylogenetic tree of the 1948 genomes based on 277533 core gene SNPs demonstrated high genetic diversity overall, with limited phylogenetic inter-mixing between isolates from humans and livestock (Fig. 1C). Pairwise SNP analysis between human- and livestock/meat-associated isolates demonstrated a median SNP distance of 41658 (range 10 - 47819, IQR 34730 - 42348), with 5 and 1 human isolates falling within 50 SNPs of livestock and meat, respectively (Fig. S5). Network analysis based on a range of SNP cut offs captured just 2 (0.1%) human isolates (from hospitals in the South East and North West) that were within 15 SNPs of livestock isolates (2 pig and 1 turkey isolate from three different farms, Fig. 2). By contrast, we observed highly related isolates (0-5 SNPs) from the same animal species on different farms (Fig. 2).

**Fig. 2.**
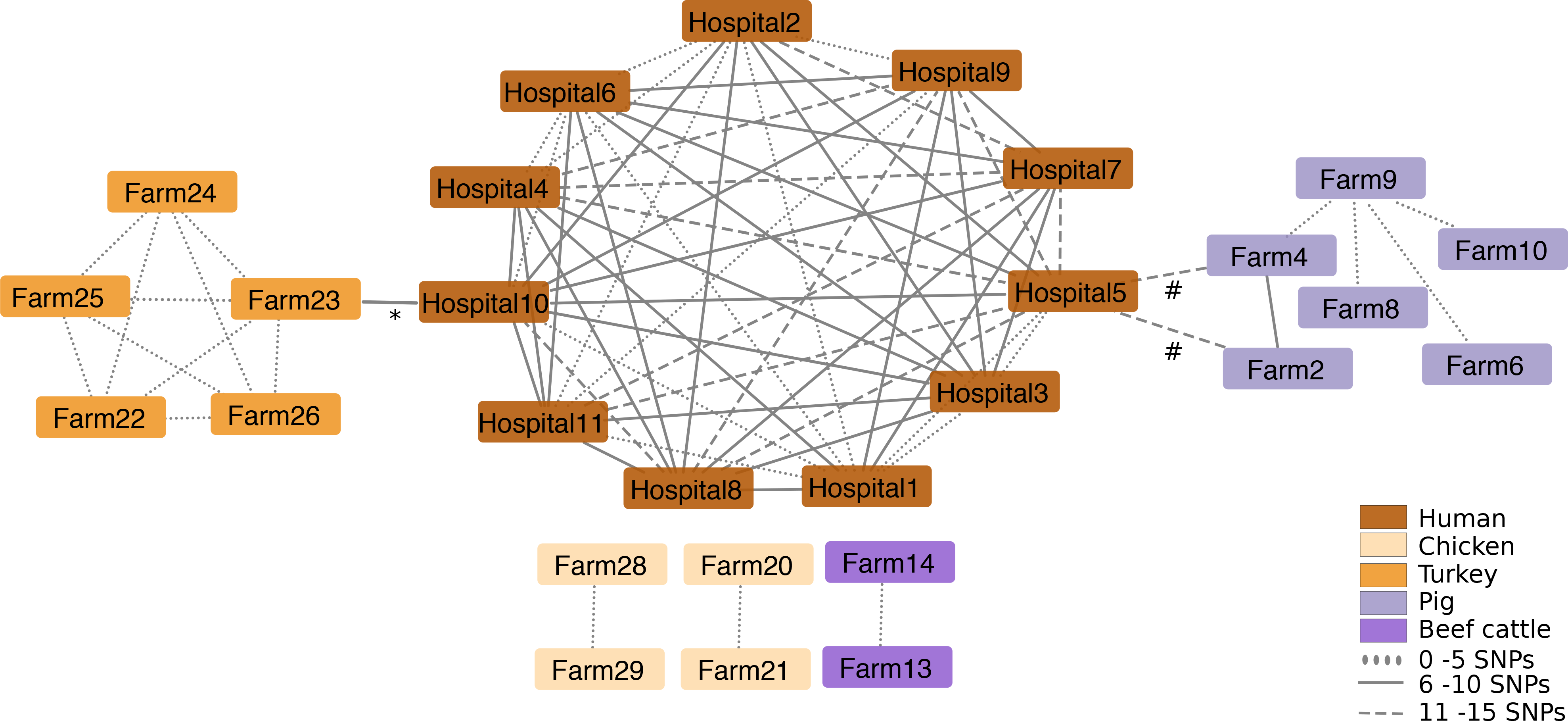
Network analysis of *E. coli* isolates cultured from livestock farms and patients with bloodstream infection isolates. The results shown are limited to those isolate pairs identified in a pairwise comparison that differed by ≤15 or less SNPs in the core genome. The place of origin for each isolate pair are connected by lines, and the style of the line reflects the SNP distance. *indicates one human isolate from hospital 10 linked to two turkey isolates from farm 23 that differed by 10 and 12 SNPs, respectively. # indicates one human isolate from hospital 5 linked to one pig isolate from farm 4 (differed by 10 SNPs) and 2 (probably duplicate) pig isolates from farm 2 that differed by 14 SNPs

We evaluated and compared the accessory (non-conserved) genome of the 1948 study isolates using principal component analysis (PCA). Principal components (PC) 1 and 2, which accounted for 50.5% and 8.3% of the variation within the data, respectively, separated the collection into two main clusters (referred to as Group 1 or Group 2). Group 1 predominantly contained human isolates and the other contained a mixture of human and livestock isolates (Fig. S6a). PCA also showed that isolates from the same STs clustered together and formed distinct sub-clusters within Group 1 and 2 (Fig. S6b). Table S3 lists the top 100 genes from PC1 and PC2 that were most strongly associated with Group 1 or 2.

### Genetic analysis of antimicrobial resistance genes and associated mobile genetic elements

Screening of the 1948 isolates for accessory genes encoding antibiotic resistance revealed that 41 different resistance genes were present in isolates from both humans and livestock (Fig. 3a). The prevalence of resistance genes in the two groups varied considerably, with some predominating in one or other reservoir while others were common in both (Fig. 3b). The seven most frequently shared genes (each present in >300 isolates) conferred resistance to beta-lactams (*bla*_TEM-1_=882), sulphonamides (*sul2*=530, *sul1*=522), aminoglycosides (*strA*=509, *strB*=478), and tetracyclines (*tetA*=423, *tetB*=335). The predominant genes conferring resistance to extended-spectrum cephalosporins were *bla*_CTX-M-15_ (human=87, livestock=32) and *bla*_CTX-M-1_ (human=1, livestock=82, meat=13). No carbapenemase or colistin resistance genes were detected.

**Fig. 3.**
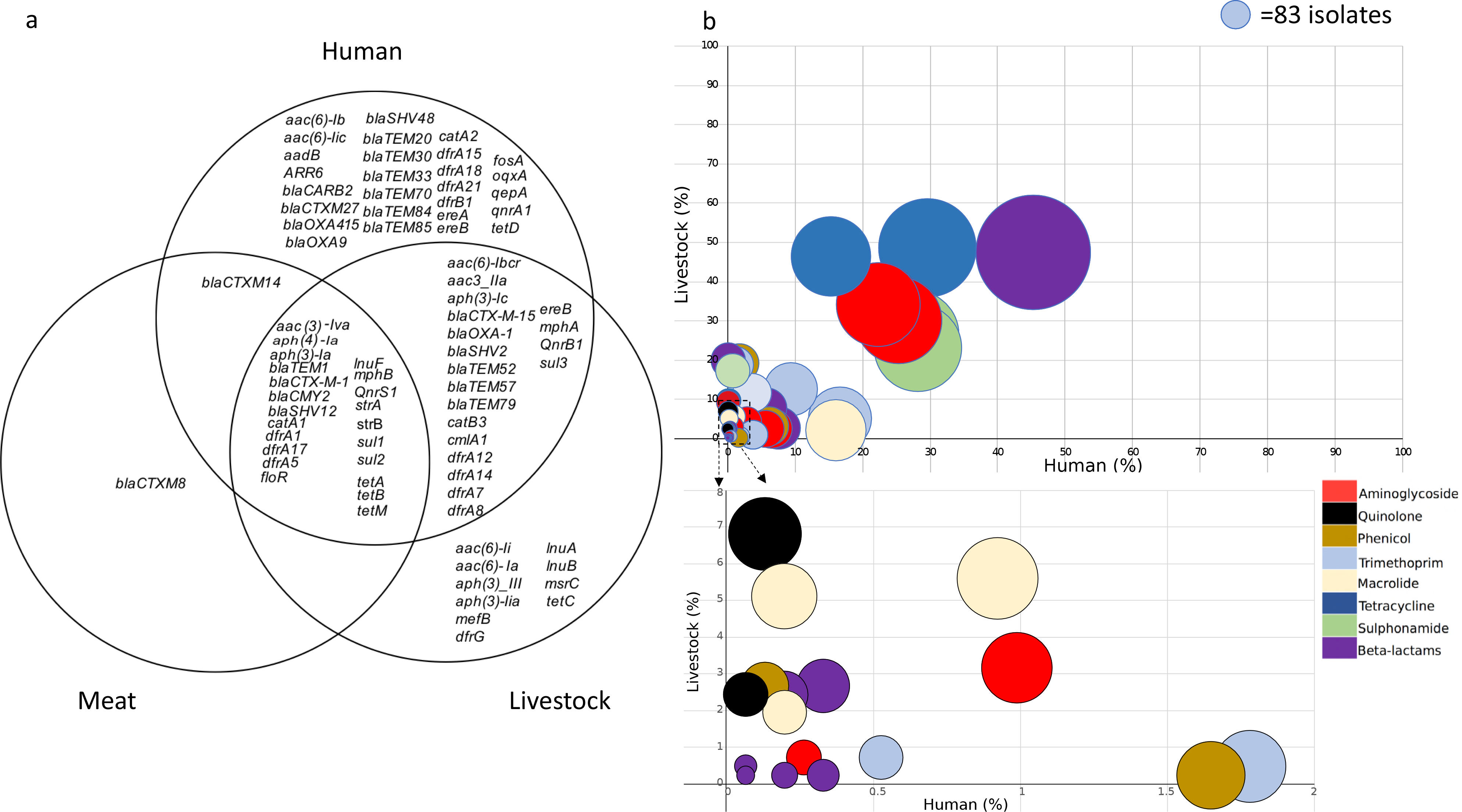
a: Venn diagram displaying antibiotic resistance genes identified in 1948 *E. coli* isolates cultured from livestock and patients with bloodstream infection. b: Bubble graph showing the proportion of genes shared between *E. coli* from humans and livestock. Lower graph shows an expanded view of very low prevalence genes that are clustered in the lower left-hand corner of the graph. The size of each bubble represents the number of isolates that the gene was identified in. Bubbles are coloured by antibiotic class

To better understand whether genes encoding resistance in isolates from livestock and humans were carried by the same or different mobile genetic elements, contigs containing the specific gene of interest were extracted and isolates clustered using hierarchical cluster analysis based on contig presence/absence. This was performed for each of the 7 most common resistance genes (*bla*_TEM-1_, *sul2*, *sul1*, *strA*, *strB*, *tetA* and *tetB*) and the 2 most prevalent ESBL genes (*bla*_CTX-M-15_ and *bla*_CTX-M-1_). Using a height cut-off value of 0 (corresponding to an identical MGE contig carriage profile), clusters were screened for presence of isolates derived from both human and livestock/meat. The majority of human isolates did not reside in clusters containing animal samples, the exception being *bla*_CTX-M-1_ where a single human isolate carrying this gene clustered with 23 animal isolates (Fig. 4). For the nine resistance genes analysed, between 0.6% and 9.8% of human isolates carrying a resistance gene shared a cluster with livestock isolates. The lowest frequency of relatedness was observed for *bla*_TEM-1_ (3 clusters, involving isolates from humans=4, livestock=3), followed by *sul1* (3 clusters, humans=3, livestock=8), *strA* (6 clusters, humans=11, livestock=22), *tetA* (6 clusters, humans=7, livestock=19), *tetB* (5 clusters, humans=11, livestock=9), *bla*_CTX-M-15_ (1 cluster, humans=5, livestock=10) *sul2* (9 clusters, humans=41, livestock=38, meat=1), *strB* (8 clusters, humans=33, livestock=29) and *bla*_CTX-M-1_ (1 clusters, humans=1, livestock=17, meat=6) (Table S4). Individual clusters often contained different STs (Table S4), which is indicative of horizontal transfer of mobile genetic elements between lineages.

**Fig. 4.**
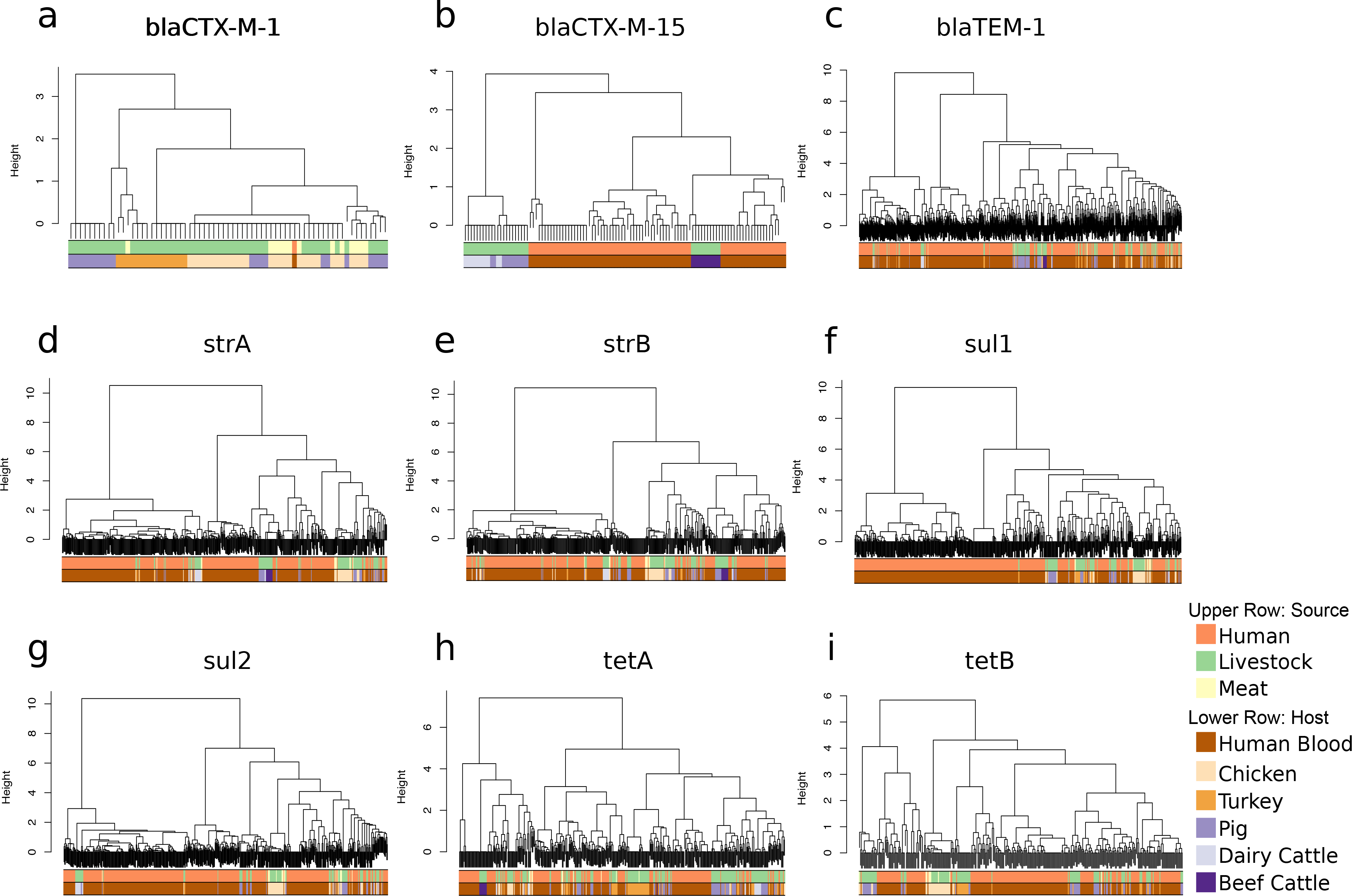
Dendrograms of mobile genetic element clusters identified for *bla*_CTX-M-1_ (a), *bla*_CTX-M-15_ (b), *bla*_TEM-1_ (c), *strA* (d), *strB* (e), *sul1* (f), *sul2* (g), *tetA* (h), and *tetB* (i) in livestock, humans and retail meat

Further characterisation of *bla*_CTX-M_ plasmids was undertaken using long-read sequencing. Two livestock-human isolate pairs positive for *bla*_CTX-M-1_ or *bla*_CTX-M-15_, respectively, were selected for sequencing using the PacBio instrument. Illumina reads for the entire study collection were then mapped to the complete plasmid assemblies of these four isolates. The single *bla*_CTX-M-1_ positive human isolate contained an IncI1 *bla*_CTX-M-1_ plasmid that was highly similar (>99% identity and ≥98% coverage) to 28 livestock isolates (chicken=18, chicken meat=8, pig=2) belonging to four different STs. By contrast, the *bla*_CTX-M-15_ plasmid in the livestock (E01) and human (D01) isolate pair were dissimilar (17% sequence shared at 99% ID) and had different replicon types (E01=IncHI2, D01=IncFIA and FII fusion). The human *bla*_CTX-M-15_ plasmid (D01) was not identified in any other isolate (human or livestock), while the livestock *bla*_CTX-M-15_ plasmid (E01) was found in other livestock isolates from the same pooled faecal sample from 1 beef farm.

We then investigated whether *bla*_CTX-M-15_ could be shared on a smaller transposable element. A 7926bp region encoding *bla*_CTX-M-15_ that was identical to a Tn*3* transposon previously identified from *E. coli* plasmid GU371928 (21) was detected in 22/32 (69%) livestock isolates (pig=11, dairy cattle=11) from 4 farms, and 3/87 (3%) of *bla*_CTX-M-15_ - positive human isolates from 2 hospitals, one of which was located in the East of England. This 7926bp region was flanked by 5bp direct repeats of TTTTA, indicating its potential for transfer between isolates.

### Plasmid-associated virulence genes

We detected a 10701 bp region in the human isolate *bla*_CTX-M-15_ plasmid (D01) that was ≥99% identical and had ≥99% coverage to a previously described virulence cassette found on large *E. coli* virulence plasmids associated with acute cystitis and neonatal meningitis (22, 23). This region contained genes encoding virulence and fitness traits including enterotoxicity, iron acquisition and copper tolerance. Screening all livestock and human isolates for this cassette showed that 28% of human invasive isolates carried a highly similar element with ≥99% coverage against the 10701 bp region and all had ≥99% sequence identity against the region covered. This virulence cassette was absent in all livestock isolates. Carriage of this element was found in isolates from all 11 hospitals, in both ESBL-positive (8%) and ESBL-negative isolates (24%), and in 51 STs.

## DISCUSSION

We investigated the prevalence and genetic relatedness of *E. coli* from livestock, meat and humans in a highly defined geographical region using a ‘One Health’ approach. ESBL-*E. coli* was isolated from 55% of livestock farms, with a frequency of ESBL-*E. coli* in different livestock species that was consistent with previous findings (24). In addition, ESBL-*E. coli* was found in 20% of pre-packaged fresh meat products. The high prevalence of ESBL-*E. coli* in chicken meat (16/30, 53%) is similar to previous studies conducted in the UK and the Netherlands (12, 25, 26). However, *E. coli* from livestock were not closely related to isolates causing human disease in our region, suggesting that livestock are not a direct source of infecting isolates and that human invasive *E. coli* are not being shared with livestock. By contrast, highly related isolates were identified between the same livestock species on different farms. Previous studies in the Netherlands that compared isolates from clinical and livestock sources using MLST indicated that the same ST could be isolated from humans and livestock (12–14, 27). We replicated this finding for CC10 and CC117, but using the more discriminatory sequence-based analysis identified that isolates from the two reservoirs were genetically distinct. A study of cephalosporin-resistant *E. coli* in the Netherlands (15) reported genetic heterogeneity between human and poultry-associated isolates but closely related isolates from farmers and their pigs. Here, we included ESBL-positive and non-ESBL *E. coli*, an important feature of the study since the majority of *E. coli* human infections in the UK are due to non-ESBL *E. coli* (16).

Screening of *E. coli* isolates from livestock, meat and humans with serious infection revealed the frequency of antimicrobial resistant genes in each reservoir and confirmed the presence of similar antimicrobial resistance genes in both livestock and humans, including *bla*_TEM-1_, *sul2*, *sul1*, *strA*, *strB*, *tetA*, *tetB, bla*_CTX-M-15_ and *bla*_CTX-M-1_. These genes confer resistance to four antibiotic classes, all of which are used in both livestock and humans (28). This confirms their ubiquitous distribution but does not provide evidence for recent transfer of genes between the two reservoirs. To address this, we hypothesised that recent sharing would be associated with transmission via the same or highly related mobile genetic elements (MGEs), as previously suggested for ESBL genes (15). Previous studies have highlighted the challenge in reconstructing plasmids and other mobile elements encoding resistance genes from whole genome sequencing (29, 30), hindering our understanding of the transmission dynamics of resistance genes. We developed an approach to detect and genetically compare mobile elements across our large study collection, with validation of findings for ESBLs using long-read sequencing. The findings from this were consistent with predominantly distinct mobile elements between livestock and humans, with an estimated 69/1517 (5%) human isolates potentially sharing closely related antimicrobial resistance-associated mobile elements with those found in livestock. Plasmid analysis led to the identification of a virulence cassette associated previously with cystitis and neonatal meningitis (22, 23), which was carried by a large multi-drug resistance IncFIA and FII plasmid. Screening of all study isolates demonstrated that this was only found in human isolates.

A limitation of our study is that we did not include all possible sources of *E. coli* for humans (for example, vegetables, fruits and pets), although a recent study found no *E. coli* with CTX-M-15 (the dominant human ESBL type) in retail meat, fruit and vegetables in five UK regions (25). We acknowledge that the *E. coli* from humans pre-dated the surveys of farms and retail meat, but we accounted for this by identifying relatedness based on a 0-15 SNP cut-off given the estimated *E. coli* mutation rate of 1 SNP/core genome/year (18, 19).

In conclusion, this study has not generated evidence to indicate that *E. coli* causing severe human infection in our region were derived from livestock, with host-specific *E. coli* lineages identified from hospitals versus farms. We identified limited sharing of antimicrobial resistance genes between livestock and humans based on long-read sequencing and analysis of mobile genetic elements. In addition, we found a known virulence cassette to be specific to human invasive samples. Further investigations are required to pursue the identification of the source of *E. coli* and resistance genes in isolates associated with severe human disease.

## METHODS

### Sampling of livestock faeces and retail meat

A cross-sectional survey was performed between August 2014 and April 2015 to isolate *E. coli* at 20 livestock farms (10 cattle & 10 pig) in the East of England. A pooled sample of approximately 50g of freshly passed faecal material was collected from each major area in a given farm (such as different pens) using a sterile scoop (Sterilin™ X400, Thermo Fisher Scientific, Loughborough, United Kingdom). Each pool was placed into a dry sterile 150ml container (Sterilin™ Polystyrene Containers, Fisher Scientific). A median of 4 samples (range 1-5) were taken from each cattle farm, and a median of 4.5 samples (range 3-9) were taken from each pig farm, resulting in a total of 85 pooled samples (34 cattle and 51 pig). In addition, caecal contents were collected from 2 deceased pigs at the time of necropsy.

Poultry reared at nine farms (4 chicken and 5 turkey) in the East of England were sampled at two abattoirs between February and April 2015. Two sample types were taken for each farm: (i) pooled faeces with a total weight of approximately 50 g from 10-20 transportation crates immediately after the livestock were removed; (ii) pools of caecal material from up to 10 birds after slaughter. Each sample was taken using a sterile scoop and a sterile surgical scalpel was used for each caecal dissection. A median of 4 (range 2-4) caecal pools and 4 (range 3-4) faecal pools were collected from livestock from each chicken farm, and a median of 1.5 (range 1-2) caecal pools and 2.5 (range 2-3) faecal pools were collected from livestock from each turkey farm. This resulted in a total of 49 pooled samples (29 chicken and 20 turkey). All samples were immediately refrigerated at 4°C upon return to the laboratory and processed on the same day.

In April 2015, 97 retail meat samples (beef 15, chicken 30, pork 42, turkey 7, venison 1, mixed minced pork and beef 1) were purchased from 11 supermarkets in Cambridge, UK, with 5-16 meat products collected from each supermarket that were selected to capture diversity in the products available. The country of origin for each meat product was recorded and where multiple countries/regions were stated on the packaging, all names were recorded.

### Microbiology

Pooled faecal samples were diluted 1:1 with sterile phosphate-buffered saline, mixed vigorously, and 100 μl plated onto Chromocult® Coliform Agar (VWR, Leuven, Belgium) and *Brilliance*™ ESBL Agar (Oxoid, Basingstoke, UK), which are selective chromogenic agars that support the growth of coliforms and ESBL-producing organisms, respectively. Agar plates were incubated at 37 °C for 48 hours in air prior to inspection. Enrichment cultures were also used to detect ESBL-producing *E. coli* by adding 1 ml of faecal preparation to 9 ml of tryptic soy broth containing 20 ¼g cefpodoxime and incubating for 24 hours in a shaking incubator (150 rpm) at 37°C in air, before 100 μl was plated onto *Brilliance*™ ESBL Agar and incubated for 48 hours in air. Up to 32 colonies of presumptive *E. coli* based on colony morphology were picked from samples taken on each farm.

Preparation and culture of meat samples followed the European standard ISO 6887– 2:2003. All exterior packaging was disinfected with alcohol prior to removal of meat. A 5g sample of meat was aseptically removed, added to 45 ml peptone broth and homogenised using a Stomacher® paddle blender (Stomacher®80 Laboratory System, Seward Ltd, UK) for two minutes. Samples were transferred into 50 ml Falcon™ tubes and incubated in a shaking incubator for 24 hours at 150 rpm at 37°C. After incubation, all samples, were plated onto *Brilliance*™ ESBL agar and incubated at 37°C for 48 hours. In addition, swabs were obtained from whole chicken carcasses and incubated in 3 ml brain heart infusion (BHI) broth (FlOQSwabs™, Copan Italia spa, Brescia, Italy) in a shaking incubator for 24 hours at 150 rpm at 37°C. Following incubation, 100μL was plated onto *Brilliance*™ ESBL agar and incubated as before. One colony of presumptive ESBL *E. coli* was picked from each *Brilliance*™ ESBL agar for further evaluation, with the exception of two meat samples where two colonies were picked to represent each of two distinct colony morphologies.

All bacterial colonies suspected to be *E. coli* were speciated using Matrix Assisted Laser Desorption Ionization – Time of Flight Mass Spectrometry (Bruker Daltonik, Bremen, Germany). Antimicrobial susceptibility was defined for each *E. coli* colony using the VITEK®2 system (bioMérieux, Marcy l’Etoile, France) with the AST-N206 card and calibrated against EUCAST breakpoints (http://www.eucast.org/clinical_breakpoints/).

### DNA sequencing

Bacterial genomic DNA was extracted using the QIAxtractor (Qiagen, Valencia, CA, USA) according to the manufacturer’s instructions. Library preparation was conducted according to the Illumina protocol and sequenced on an Illumina HiSeq2000 (Illumina, San Diego, CA, USA) with 100-cycle paired-end runs. Sequence data were retrieved for a further 1517 open access *E. coli* isolates associated with bloodstream infection (16, 17), 424 were isolated between January 2006 and December 2012 at the Cambridge University Hospitals NHS Foundation Trust and 1093 were submitted to the British Society for Antimicrobial Chemotherapy Bacteraemia Resistance Surveillance Project by 11 UK hospitals between 2001 and 2011 (for details, see www.bsacsurv.org, Table S1) (31). Previous description and analysis of these genomes (16, 17) did not include comparisons with isolates from livestock or meat.

### Genome Assembly, Annotation and Multilocus Sequence Typing

Taxonomic identity was assigned using Kraken (32). One isolate (VREC0294) was found to not be *E. coli* and was excluded from further analysis. *de novo* assembly of short read data was performed using Velvet (33) and assemblies were annotated using Prokka (34). Multilocus sequence types (MLST) were identified using the MLST sequence archive (https://enterobase.warwick.ac.uk). Sequence types (STs) were classified into clonal complexes using the eBURST V3 algorithm (http://eburst.mlst.net/).

### Pan-genome analysis

The pan-genome was calculated for all 1948 isolates using Roary (35), with a 90% ID cut-off and genes classified as ‘core’ if they were present in at least 99% of isolates. A maximum-likelihood tree was created using RAxML (36) based on SNPs in the core genes. Principal component analysis was performed across the 1948 isolates based on the accessory genes from Roary using R. A Spearman rho correlation analysis was performed on the principal components, the gene absence/presence data and the ST and source of isolation metadata.

### Phylogeny based analysis of individual lineages

Lineage specific-analyses were performed by mapping the sequence reads for isolates belonging to clonal complex (CC) 10 and CC117 to an *E. coli* reference genome from the same clonal complex using SMALT 0.7.4 (http://www.sanger.ac.uk/resources/software/smalt/). MG1655 K12 (ENA accession number U00096.2) was used as the reference genome for CC10, and a *de novo* assembly of the CC117 study isolate with the lowest number of contigs was used as the reference genome for CC117 as no reference genomes were available. To create a ‘core’ genome, mobile genetic elements (MGEs) were identified using gene annotation, PHAST (phast.ishartlab.com), and BLAST (https://blast.ncbi.nlm.nih.gov) and removed, together with contigs less than 500 bp in length. Recombination was removed using Gubbins (37). A maximum-likelihood phylogeny was created using RAxML (36) with 100 bootstraps and a midpoint root. Genetic diversity was calculated based on pairwise differences in SNPs in the core genomes using an in-house script. Visualisation of phylogenetic trees was performed using iToL (http://itol.embl.de) (38) and FigTree v 1.4.2 (http://tree.bio.ed.ac.uk/software/figtree/).

### Bayesian Evolutionary Analysis Sampling Trees (BEAST)

All published genomes for CC117 in the *Enterobase* online database (https://enterobase.warwick.ac.uk, accessed 10 June 2016) that had been generated on an Illumina instrument and had the country, year and source of isolation available were identified (ERR769196, ERR769195, ERR769183, ERR769169, SRR1314275, SRR3410778, SRR3438297). These were mapped to the CC117 reference using SMALT, combined with the CC117 study isolates, and mobile elements and recombination were removed as before. Dating of this lineage was completed using BEAST v.1-8 (39) BEAST v.1-8 was run using the Hasegawa, Kishino and Yano (HKY) and gamma substitution model. We compared combinations of three population size change models (constant, exponential and Bayesian skyline plot) and three molecular clock models (strict, exponential and uncorrelated lognormal). A Bayesian skyline population model and an uncorrelated lognormal molecular clock were selected based on Bayes factors calculated from path sampling and stepping stone sampling (39, 40).

### Detection of antimicrobial resistance and mobile elements

Acquired genes encoding antibiotic resistance were identified using Antibiotic Resistance Identification By Assembly (ARIBA), comparing the study genomes against an in-house curated version of the Resfinder database (41–43) consisting of 2015 known resistance gene variants. Genes were classified as present using an identity of 90% nucleotide similarity. Genes reported as fragmented, partial or interrupted were excluded.

For all *bla*_CTX-M-1_, *bla*_CTX-M-15_, *bla*_TEM-1_, *sul1*, *strA*, *strB*, *sul2*, *tetA* and *tetB* positive isolates, whole genome assemblies were screened to identify the contig carrying the AMR gene using the blastn application (44) with the AMR gene sequence as the ‘query sequence’. The identified contigs were then aligned against a previously curated database of complete Enterobacteriaceae plasmids (45) in order to filter out sequences representing *E. coli* chromosome fragments. For each AMR gene, a database containing unique AMR-carrying contigs was created using cd-hit-est (46) based on 90% identity cut-off. To determine contig carriage, for each of the eight AMR genes all isolates positive for that gene were mapped against the respective gene-specific database of contigs using Short Read Sequencing Typing (SRST2) using a minimum 90% coverage cut-off. The contig presence/absence data were converted into a distance matrix and hierarchical clustering was performed using R function *hclust* and the ward.D2 method.

To examine plasmids encoding the ESBL genes *bla*_CTX-M-15_ and *bla*_CTX-M-1_, two pairs of livestock and human isolates were selected for sequencing on the PacBio RSII Instrument (Pacific Biosciences, Menlo Park, CA, USA) (n=4) and *in silico* PCR was used to perform plasmid incompatibility group/replicon typing (47). These two genes were selected as they were the most prevalent ESBL genes found in the 1948 isolates. The pair of *bla*_CTX-M-15_ positive isolates was selected as they contained 5 identical genes encoding antimicrobial resistance. The single human *bla*_CTX-M-1_ positive isolate in the collection was selected and a livestock isolate with the most similar resistance gene profile was selected, with both isolates containing 3 identical genes encoding antimicrobial resistance. DNA was extracted using the phenol/chloroform method (48) and sequenced using the PacBio RS II instrument. Sequence reads were assembled using HGAP v3 (49) of the SMRT analysis software v2.3.0 (https://github.com/PacificBiosciences/SMRT-Analysis), circularized using Circlator v1.1.3 (50) and Minimus 2 (51) and polished using the PacBio RS_Resequencing protocol and Quiver v1 (https://github.com/PacificBiosciences/SMRT-Analysis). Fully assembled plasmids were compared using WebACT (http://www.webact.org) and BLASTn (https://blast.ncbi.nlm.nih.gov).

### Ethical approval

The study protocol was approved by the CUH Research and Development Department (ref: A093285) and the National Research Ethics Service East of England Ethics Committee (ref: 12/EE/0439 and 14/EE/1123).

## DATA ACCESS

Sequence data for all isolates have been submitted to the European Nucleotide Archive (www.ebi.ac.uk/ena) under study accession number PRJEB4681 (all human *E. coli*), PRJEB8774 (non-ESBL-producing *E. coli* from livestock) and PRJEB8776 (ESBL-producing *E. coli* from livestock and meat), with the accession numbers for individual isolates listed in Table S1.

## AKNOWLEDGMENTS

The authors thank the Wellcome Trust Sanger Institute core library construction, sequence and informatics teams, and the Pathogen Informatics team. We thank the staff at farms and abattoirs for assistance in sample collection and Elizabeth Lay for laboratory support during the meat survey. We thank Olivier Restif for statistical advice. The flocked swabs used in the meat survey were donated by Copan Italia spa. This publication presents independent research supported by the Health Innovation Challenge Fund (WT098600, HICF-T5-342), a parallel funding partnership between the Department of Health and Wellcome Trust. The views expressed in this publication are those of the author(s) and not necessarily those of the Department of Health or Wellcome Trust. This project was also funded by a grant awarded to the Wellcome Trust Sanger Institute (098051). TG is a Wellcome Trust Research Training Fellow (103387/Z/13/Z). CL is a Wellcome Trust Sir Henry Postdoctoral Fellow (110243/Z/15/Z). FC is a Wellcome Trust Sir Henry Postdoctoral Fellow (201344/Z/16/Z). DJ and JP are funded by the Wellcome Trust grant 098051 M.dG is funded by the Medical Research Council (MR/K021133/1).

## AUTHOR CONTRIBUTIONS

Study design: CL, TG and SJP. Developed study protocols: CL and TG. Collection of farm samples: CL and TG. Sourcing and microbiological culture of retail meat samples: NH, TG and MH. Obtained access to farms and abattoirs: JHG, PW, MR and MH. Bacterial identification, susceptibility testing and phenotypic testing: CL, BB and PN. Bioinformatics analyses and interpretation: CL, DJ, KER, FC and MdG. Figure production: CL, DJ, FC and MdG. Contributing sequence data: NMB and CH. Writing the manuscript: CL and SJP. Responsibility for supervision and management of the study: JP and SJP. All authors read and approved the final manuscript.

## SUPPLEMENTAL MATERIAL

The following Supplemental material is available for this article:

**Fig. S1.** Maximum likelihood phylogenetic tree of the 1517 invasive isolates from 11 UK hospitals isolated from 2001-2012. The tree annotated by ‘association’ (community, healthcare or unknown).

**Fig. S2.** Maximum likelihood tree of isolates from animals and human invasive infections from Clonal Complex 10 based on SNPs in the core genes.

**Fig. S3.** Maximum likelihood tree of the isolates from animals and human invasive infections from Clonal Complex 117 based on SNPs in the core genes.

**Fig. S4.** Dated Bayesian maximum likelihood tree of CC117 isolates from animals and human invasive infections from the UK and 7 isolates from Denmark and the United States of America.

**Fig. S5.** Barplot of pairwise SNP differences between 1928 isolates from humans (n=1517) and livestock (n=41), with the frequency representing the number of human isolates related to a livestock isolate at the SNP threshold defined on the X axis. A range of SNPs from 0-500 is shown.

**Fig. S6.** Principal component analysis based on the presence/absence of accessory genes in animals, humans and meat. (a) shows principal component (PC) 1 (x-axis) against PC2 (y-saxis) labelled by source, (b) shows principal component (PC) 1 (x-axis) against PC2 (y-axis) labelled by major sequences types (STs).

**Table S1:** Details of all *E. coli* samples sequenced from animals, meat and humans (see excel S1)

**Table S2:** Details of meat samples collected as part of survey (see excel S2)

**Table S3:** The top 100 genes from PC1 and PC2 that are most strongly associated with either Group 1 or Group 2 (see excel S3)

**Table S4:** Details of common mobile elements (denoted as “profiles”) found in animal, humans and meat for *bla*_CTX-M-1_, *bla*_CTX-M-15_, *bla*_TEM-1_, *strA*, *strB*, *sul1*, *sul2*, *tetA* and *tetB* (see excel S4

